# Attention-based GCN Integrates Multi-omics Data for Breast Cancer Subtype Classification and Patient-specific Gene Marker Identification

**DOI:** 10.1101/2022.09.05.506572

**Authors:** Hui Guo, Xiang Lv, Yizhou Li, Menglong Li

**Author notes:** Corresponding author. (Li M), (Li Y).

## Abstract

Breast cancer is a heterogeneous disease and can be divided into several subtypes with unique prognostic and molecular characteristics. Classification of breast cancer subtypes plays an important role in the precision treatment and prognosis of breast cancer. Benefitting from the relation-aware ability of graph convolution network (GCN), we presented a multi-omics integrative method, attention-based GCN, for breast cancer molecular subtype classification using mRNA expression, copy number variation, and DNA methylation multi-omics data. Several attention mechanisms were performed and all exhibited effectiveness in integrating heterogeneous data. Column-wise attention-based GCN outperformed the other baseline methods, achieving AUC of 0.9816, ACC of 0.8743 and MCC of 0.8151 in 5-fold cross validation. Besides, Layer-wise Relevance Propagation (LRP) algorithm was used for interpretation of model decision and could identify patient-specific important biomarkers which were reported to be related to the occurrence and development of breast cancer. Our results highlighted the effectiveness of GCN and attention mechanisms in multi-omics integrative analysis and implement of LRP algorithm could provide biologically reasonable insights into model decision.

**Author Summary:** Identification of molecular subtype is essential to understanding the pathogenesis of cancer and advancing the development of cancer precision treatment. We have developed a graph convolution network architecture to predict the molecular subtype of breast cancer patients based on multi-omics data. The major difference between our work and other methods is that we have offered a new view for multi-omics data integration with the assistance of graph convolution network. Besides, we adopted an interpretability method to explain the model decision and prioritize the genes for biomarker discovery.

## Introduction

Breast cancer is highly heterogeneous showing different clinical behavior and treatment responses. At present, the classification of breast cancer based on pathologic features and IHC evaluations has been well established for clinical application and validity. The latest AJCC-TNM 8 staging not only uses clinical and pathologic information but also incorporates immunophenotype-related prognostic markers (ER/PR/HER2) into the prognostic staging. High-throughput sequencing technologies can reveal the heterogeneity of breast cancer at the molecular level. Genome-wide expression profiling studies classified breast cancer into five intrinsic subtypes, namely luminal A(LumA), luminal B(LumB), HER2-overexpressing(Her2), basal-like (Basal), and normal-like(Normal) tumors[1]. At the same time, there is a large correlation between breast cancer intrinsic subtypes and immune phenotypes, and different cohort studies have reproduced similar intrinsic subtypes by using different numbers of gene signatures[2,3]. It also indicates that multiple genes act synergistically as functional modules leading to the inherent molecular heterogeneity of breast cancer. Multi-omics sequencing platform offers opportunities to study breast cancer at various molecular levels[4]. Each type of omics data exhibits specific disease associations and several studies have developed machine learning models for breast cancer subtype identification and also exploit subtype-specific risk predictors or biomarkers[5–8].

However, single omics data analysis is limited to correlations, mainly reflecting reactive processes rather causative ones. Integrating different omics data may provide more biological insights[9], for example, further refining the intrinsic classification of breast cancer(intCluster10[10]) or improving the predictive power of diagnosis model[11].

Many multi-omics integrated analysis approaches are based on machine learning techniques, aiming at patient stratification[12–14], biomarker discovery[15,16], pathway analysis[17], and drug discovery[18]. One of the challenges of multi-omics analysis is the high dimensionality of data and many studies have proposed different deep learning integration strategies[19–21]. Autoencoders are designed to reconstruct the original high-dimensional input into latent low-dimensional features which can represent the raw data and have been successfully applied to analyze high-dimensional omics data and integrate heterogeneous data types[22]. Generative adversarial networks (GAN) can also be used to extract effective feature representations from multi-omics heterogeneous[23]. Network-based fusion strategies were also applied in multi-omics data integration[24]. The application of GCN in multi-omics integration analysis is seldomly reported. And most of the existing studies usually combine feature representation learning technology with GCN based on the patient similarity network for patient diagnosis or clustering[25,26], where the prior structural relationships between genes are not taken into account. Genes act together in signaling, regulatory pathway, and protein complexes. The information contained in the protein–protein interaction network (PPI) is highly important for gene function prediction[27], gene module detection[17], and novel cancer gene discovering[28,29]. Here, we developed a GCN based framework that could simultaneously incorporate multi-omics features of genes and their structural relationships with gene-gene interaction network as graph input.

Genomic changes have complicated associations with cancer phenotypes, and each type of molecular features is not independent. For example, CNV causing a gene expression change may be relevant to disease, and presence of DNA methylation near the transcription start site of a gene is associated with its expression change. Most of existing supervised multi-omics can be divided into two types, concatenation-based methods and ensemble-based methods. Concatenation-based methods integrate different types of data input by directly concatenating the data input features or the learned embedding features, and the latter is implemented by integrating predictions from different classifiers. Both of them fail to catch the cross-omics correlations and are likely to be biased towards certain omics data types considering the existence of systematic noises or scale differences[25,30]. Several supervised machine learning classification methods are developed for modeling the cross-omics correlations. For example, Data Integration Analysis for Biomarker discovery using Latent components (DIABLO) maximizes the common or correlated information between multiple omics datasets and discriminates phenotypic groups[31]. A sparse multiple discriminative canonical correlation analysis (SMDCCA) integrates differentially expressed genes (DEGs), differentially methylated CpG sites (DMCs), and differential metabolic products (DMPs) for osteoporosis risk and identifies biomarkers which are correlated spanning different biological layers[32]. Attention mechanism is one of the most important concepts in the deep learning field and can explain the incomprehensible network architecture behavior[33]. We introduced attention-based GCN to capture the complex correlations between different omics date types.

The lack of transparency is a significant disadvantage of *black-box* deep learning methods, limiting the interpretability and thus the scope of application in practice[34]. In cancer diagnostics, the demand for model interpretation is increasing since it’s crucial to identify important biomarkers and related biological processes involved in decision making process and further convince physicians in model rationality. What’s more, reliable interpretation strategies can help investigate biological meanings of the nonlinear deep models and explore biological patterns which may lead to new biological hypotheses. Layer-wise Relevance Propagation (LRP) algorithm may provide a better explanation of model decision than the sensitivity-based approach or the deconvolution method in image classification[35]. LRP has been applied in explaining the deep neural network for gene expression[36] and to visualize convolutional neural network decisions for Alzheimer’s disease based om magnetic resonance imaging data.

In summary, to explicitly model the associations between different molecular-level features of genes and their associations with disease phenotypes, we mapped features at multiple levels (mRNA, DNA methylation and CNV) to the gene level and adopted GCN with prior structural information from protein-protein interaction network as graph input for feature fusion. Based on multi-omics data from TCGA BRCA, we proposed an attention-based GCN for classification of different breast cancer molecular subtypes and jointly learning associations across multiple molecular levels of genes. We compared the model performance of GCN with three types of attention mechanism approaches. The results show that GCN combined with omics-specific attention mechanism can significantly improve the prediction performance of neural network. We also analyzed the prediction performance under different molecular-level data types and proved that attention-based GCN has certain advantages in integrating high-dimensional features. Furthermore, we adopted LRP algorithm to explain model decisions and identify patient-specific gene markers and related functional modules. Finally, we validated the generalization performance of the model on unlabeled data. (**Fig 1**)

**Fig 1.**
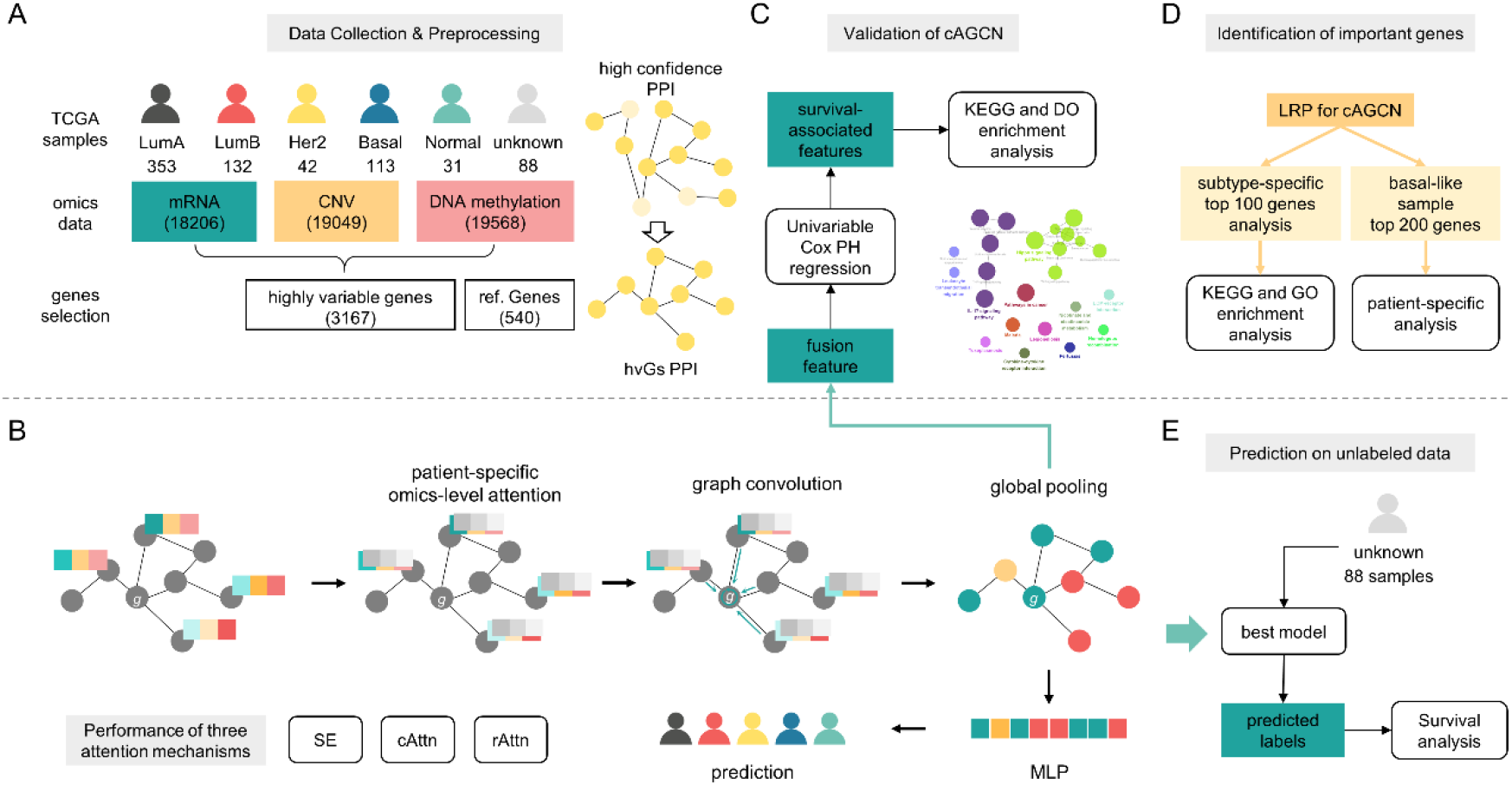
Workflow overview. **A**. Data collection and preprocessing of TCGA BRCA multi-omics data and PPI network. 67l samples with subtype information were used for model training. 3167 highly variable genes (hvGs) of each omics type were retained and 540 reference genes were downloaded from DisGenet. Raw PPI was collected from StringDB (v11.5) and edges with low confidence score (<850) were filtered out. Subnetwork consisting of hvGs was extracted from high confidence PPI with consideration of keeping different hop neighborhood relationships. **B**. Attention-based graph convolution network (AGCN) framework. AGCN takes multi-omics data and corresponding subnetwork as input. Performance of three attention mechanisms (SE, cAttn and rAttn) were discussed. **C**. Validity of AGCN. Univariable Cox PH was adopted to screen genes associated with patient overall survival based on the embedded feature extracted from global pooling layer of AGCN and raw expression data respectively. KEGG and DO enrichment analysis were performed on the significant genes. **D**. Interpretation of AGCN. Layer-wise relevance propagation algorithm was adopted on the AGCN model for feature ranking. KEGG and GO enrichment analysis were performed on top100 genes of each subtype. Specifically, patient-specific pattern of top200 genes in basal-like samples was further discussed. **E**. Validation on unknown data. The best model obtained from cross-validation was used to predict 88 unlabeled samples and further survival analysis was performed based on predicted labels.

## Results

### Performance of three attention mechanisms for multi-omics feature fusion and interpretation for attention-based GCN model

Based on the constructed hvGs dataset (hvG-3167) and corresponding hvG-PPI subnetwork, the performances of the GCN model using three different attention mechanisms and the classical MLP model framework were tested respectively. The classification results are shown in **Table 1**. We observed that the simple GCN model performed worst and adding different attention mechanisms could significantly improve the performance of the GCN model, suggesting that the introduced attention mechanisms were effective for feature fusion at different molecular levels. Additionally, cAGCN outperformed cAMLP indicating that graph convolution can further improve the representation ability of the model. The rAGCN is slightly inferior to the performance of cAGCN. The computational complexity of the attention mechanism is O(N^2^T), where N is the size of Q/K and T is the size of V. Considering the computational cost and results above, cAGCN was chosen as the best model (AUC: 0.9816±0.0084, ACC: 0.8793±0.0315, MCC: 0.8151±0.0482). Among all samples of the validation data set, cAGCN achieved a relatively higher recall and precision score on basal-like samples which accounts for 17.16% of training samples demonstrating that cAGCN is not easily affected by unbalanced data set. Normal samples are relatively more difficult to be classified accurately and more likely to be identified as luminal A samples (**Fig 2C**) which agrees with the discovery that normal-like breast cancer is similar to luminal A disease and is characterized by hormone-receptor positive (estrogen-receptor and/or progesterone-receptor positive), HER2 negative, and low levels of the protein Ki-67 while it has a slightly worse prognosis than luminal A cancer’s.

**Table 1.** Performance of the GCN model using three different attention mechanisms and the classical MLP model framework.

| Model | AUC | ACC | MCC |
| --- | --- | --- | --- |
| simple GCN | 0.8596±0.0549 | 0.6154±0.0664 | 0.3345±0.1394 |
| SEGCN | 0.9705±0.0116 | 0.8554±0.0370 | 0.7734±0.0600 |
| cAMLP | 0.9477±0.0126 | 0.7929±0.0304 | 0.6788±0.0497 |
| cAGCN | 0.9816±0.0084 | 0.8793±0.0315 | 0.8151±0.0482 |
| rAGCN | 0.9650±0.0166 | 0.8763±0.0172 | 0.8114±0.0283 |

*Note:* SEGCN: GCN+SE; cAMLP: MLP+cAttn; cAGCN: GCN+cAttn; rAGCN: GCN+rAttn

**Fig 2.**
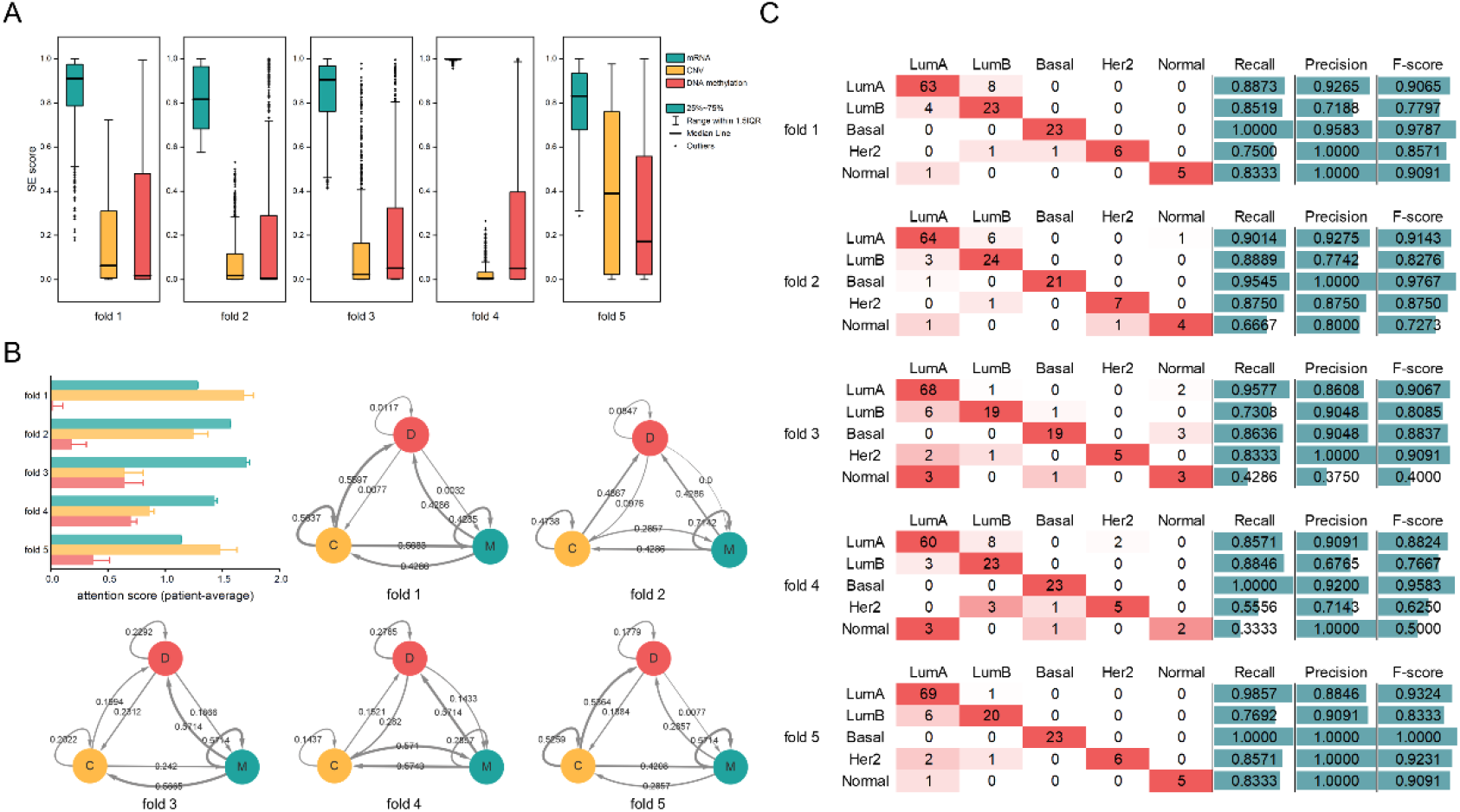
Performance of three attention mechanisms for multi-omics feature fusion. **A**. Boxplot shows distribution of SE attention score of three omics data types. **B**. Barplot (upper-left) of cAttn attention score of each omics type and ternary entity diagram of relationships between omics pairs in five datasets. M: mRNA, green. C:CNV, yellow. D: DNA methylation, red. **C**. Heatmap of confusion matrix obtained from cAGCN. Three metrics (recall, precision and F-score) were calcuated for each subtype.

The attention mechanism can explain the inherent decision-making logic of the model, thus increasing the model interpretability. Both SE and cAttn can reflect the contribution of different omics types to model decision-making. The boxplot of attention score extracted from SEGCN shows the patient-specific omics contribution of different models in 5-fold cross-validation (**Fig 2A**, Table S1). It can be observed that the information of mRNA is retained to the maximum extent, and CNV and DNA methylation data also have a certain degree of contribution, but the SE score is close to 0 in some sample decisions. We further visualized the sample-averaged attention score obtained from cAGCN with a heatmap reflecting the correlation between omics (**Fig 2B**). There are some differences in the distribution of attention scores when the data set varies. We supposed that cAGCN yielding a better prediction score is attributed to its advantage in learning the cross-omics complicated relationships from heterogeneous data. Overall, mRNA data plays an important role in the model decision. It is inferred that the molecular heterogeneity of breast cancer was largely captured at the level of gene transcription and protein function.

### Performance of the model under different hyperparameters

We mainly investigated the influence of two hyperparameters on the cAGCN model. Considering that network shrinkage would affect the quality of the input graph and then the performance of the GCN, we tested the graph input with different order neighborhoods retained (*k*-hop). As the retention order increases, the density of the input graph gets larger. And when it exceeds a certain order (k=3), the performance of the model decreases obviously (**Table 2**). Keeping higher-order neighborhood relationships may produce more noise edges and graph Laplacian with sparse networks can play a certain role as regularization[37] and improve the generalization performance of the model. The multi-head attention mechanism is an essential component of cAGCN, and the hyperparameter optimization is carried out on the number of attention heads. In the initial stage, MCC and ACC tend to increase along with the increase of the number of attention heads, and AUC is not sensitive to hyperparameters (**Fig** S1). Finally, the number of attention heads was set as 7 and the 2-hop subnetwork was considered as graph input.

**Table 2.** Results of cAGCN’s graph input with different order neighborhoods retained (*k*-hop).

| $k$-hop | hvG-3167 | | | |
| --- | --- | --- | --- | --- |
|  | AUC | ACC | MCC | Density |
| 1 | 0.9672±0.0118 | 0.8689±0.0239 | 0.7953±0.0384 | 0.0019 |
| 2 | 0.9681±0.0084 | 0.8793±0.0315 | 0.8151±0.0482 | 0.0342 |
| 3 | 0.9744±0.0059 | 0.8748±0.0323 | 0.8071±0.0486 | 0.2487 |
| 4 | 0.9679±0.0059 | 0.8300±0.0569 | 0.7312±0.0937 | 0.6142 |
| 5 | 0.9573±0.0276 | 0.8405±0.0848 | 0.7534±0.1289 | 0.8274 |

### Performance of cAGCN under different omics data types

When comparing the hvGs of three molecular level data, it can be found that there is relatively little overlap among the three gene sets (**Fig 3A**). We further explored the ability of different omics data types to predict the molecular subtype of breast cancer. Seven data sets were constructed under different permutations and combinations (**Table 3**). Classification performance of single-omics data set is significantly inferior to that of multi-omics data set. Although the number of multi-omics features has greatly increased, using proper feature fusion strategy can still achieve good classification performance.

**Fig 3.**
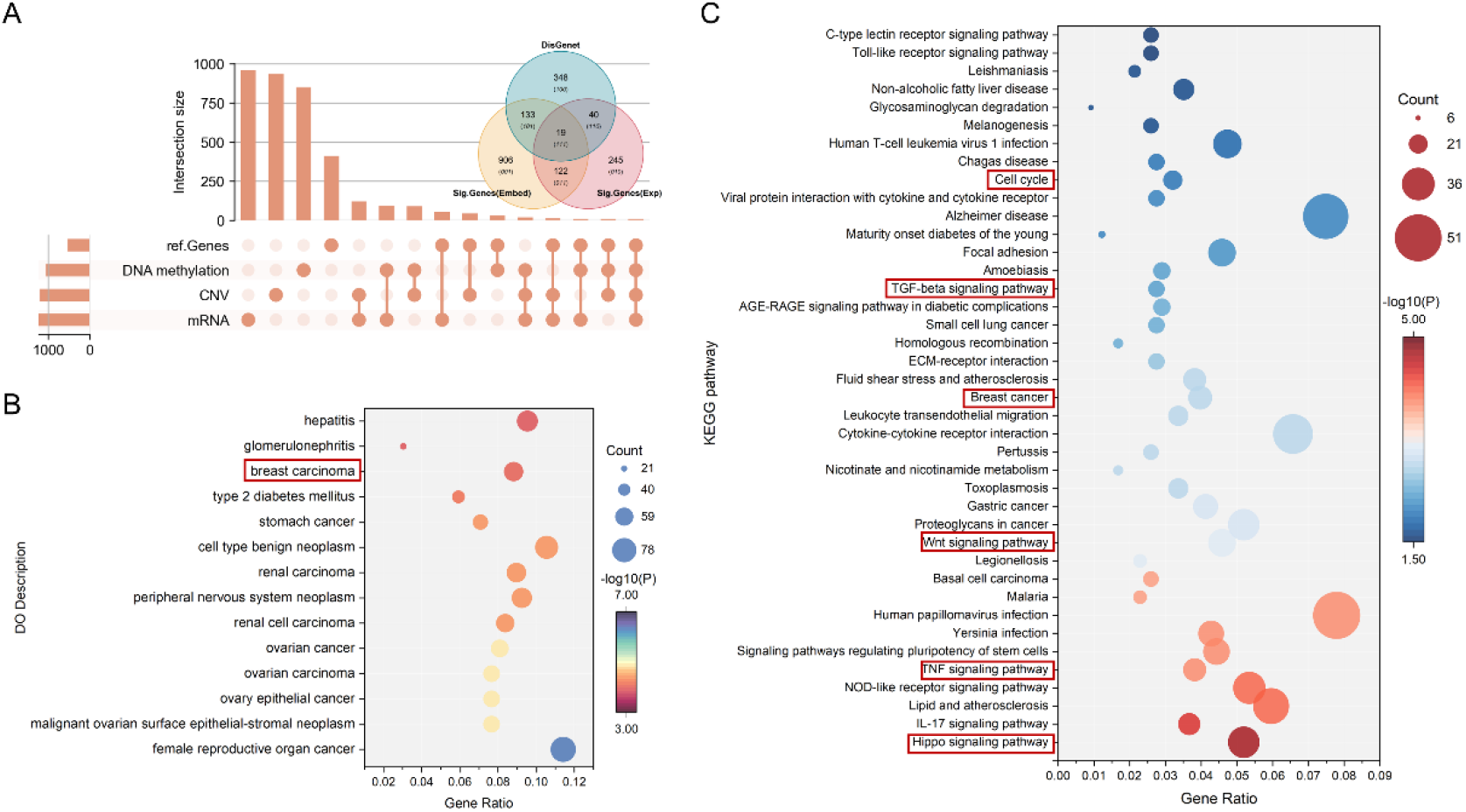
Effects of adding reference genes and enrichment analysis of significant genes. **A**. Upset plot (left) of four gene subsets and Venn plot of two significant gene sets and reference gene sets. The four gene subsets include hvGs of mRNA, CNV and DNA methylation and DisGenet reference genes (ref.Genes). Significant genes are screened based on embedded features obtained from cAGCN and raw gene expression data respectively. **B-C**. Bubble plot of DO (left) and KEGG (right) enriched terms. Bubbles are colored by P values and the size is proportional to the number of genes.

**Table 3.** Performance of cAGCN under different omics data types.

| #Dataset | num of<br>sig.genes | num of<br>features | AUC | ACC | MCC |
| --- | --- | --- | --- | --- | --- |
| M | 2000 | 1237 | 0.9169±0.1267 | 0.7139±0.0250 | 0.5349±0.0449 |
| D | 2000 | 1070 | 0.8843±0.0212 | 0.6706±0.0344 | 0.4593±0.0700 |
| C | 1974 | 1198 | 0.8459±0.0178 | 0.6185±0.0236 | 0.3464±0.0559 |
| MD | 3829 | 2200*2 | 0.9743±0.0073 | 0.8838±0.0243 | 0.8187±0.0384 |
| MC | 3762 | 2294*2 | 0.9685±0.0111 | 0.8748±0.0256 | 0.8061±0.0404 |
| DC | 3785 | 2162*2 | 0.9361±0.0253 | 0.8271±0.0186 | 0.7295±0.0307 |
| MDC | 5425 | 3176*3 | 0.9816±0.0084 | 0.8793±0.0315 | 0.8151±0.0482 |

*Note:* M: mRNA; D: DNA methylation; C: CNV

### Effects of adding reference genes and validity of cAGCN

By comparing with the reported breast cancer related oncogenes in DisGenet database, we found that the selected hvGs filtered out important breast cancer related oncogenes. Most of the hvGs have not been reported to be directly associated with breast cancer (**Fig 3A**). However, the prediction results show that these genes are closely related to the molecular subtypes of breast cancer. Meanwhile, the addition of reference gene set has little effects on the model performance (**Table 4**), indicating that these hvGs are not independent from reference genes and are probably directly or indirectly involved in the molecular processes related to the occurrence and proliferation of breast cancer.

**Table 4.** Effects of adding reference genes on model performance.

| #Dataset | num of<br>features | AUC | ACC | MCC |
| --- | --- | --- | --- | --- |
| MDC | 3176*3 | 0.9816±0.0084 | 0.8793±0.0315 | 0.8151±0.0482 |
| MDC_addRef | 3483*3 | 0.9641±0.0169 | 0.8808±0.0284 | 0.8133±0.0448 |

*Note:* MDC\_addRef: hvGs + reference genes

To further verify the effectiveness of feature fusion strategy, univariable Cox PH regression was adopted to analyze the fusion feature extracted from cAGCN middle layer. After screening out genes significantly associated with the overall survival of breast cancer, KEGG functional enrichment analysis and DO enrichment analysis were performed. The same process was carried out based on the original gene expression data. More genes are discovered to be related to the prognosis of breast cancer (**Fig 3A**) and significantly cancer-enriched. The results of DO enrichment analysis show that these genes are significantly enriched in breast cancer-related genes (**Fig 3B**). The results of KEGG pathway enrichment analysis further demonstrate the correlation between these genes and the occurrence and proliferation of cancer (**Fig 3C**). For example, Dysregulation of Hippo Signaling Pathway is directly relevant to various aspects in cancer biology, which includes cell proliferation, EMT (epithelial-to-mesenchymal transition) and metastasis[38]. TNF signaling pathway contains an important pro-inflammatory cytokine, tumor necrosis factor-α (TNF-α), which is found in the TME that is involved in all stages of breast cancer development, affecting tumor cell proliferation and survival, EMT, metastasis and recurrence[39]. Wnt signaling pathway regulates a variety of cellular processes, including cell fate, differentiation, proliferation and stem cell pluripotency. Abnormal Wnt signal is a marker of many cancers[40]. Transforming growth factor beta(TGF-β) in TGF-β signaling pathway is a multifunctional cytokine involved in regulation of cell proliferation, differentiation and survival/or apoptosis of many cells[41]. In later stage of breast cancer, TGF-β has direct pro-tumorigenic effects through the stimulation of invasion, the migration of tumor cells[42,43]. Cell Cycle disorder is a common feature of human cancer. Different breast cancer subtypes show different patterns on cell cycle and its checkpoints[4].

### Identification of important genes by LRP

We used the LRP algorithm to identify important genes that affect model decisions (Table S2). We first performed the functional analysis based on the top100 genes related to each breast cancer subtype. Although there is little genetic overlap among the subtypes, the biological processes involved in them have a high degree of functional overlap (**Fig 4A**). The results of GO and KEGG functional enrichment analysis show that the most important genes are related to cell cycle (**Fig 4B, C**, Table S3). Based on the merged network, 12 MCODE components are identified, where MCODE 2 represents mitotic cell cycle and MCODE 11 is related to Wnt signaling pathway (**Fig 4D**).

**Fig 4.**
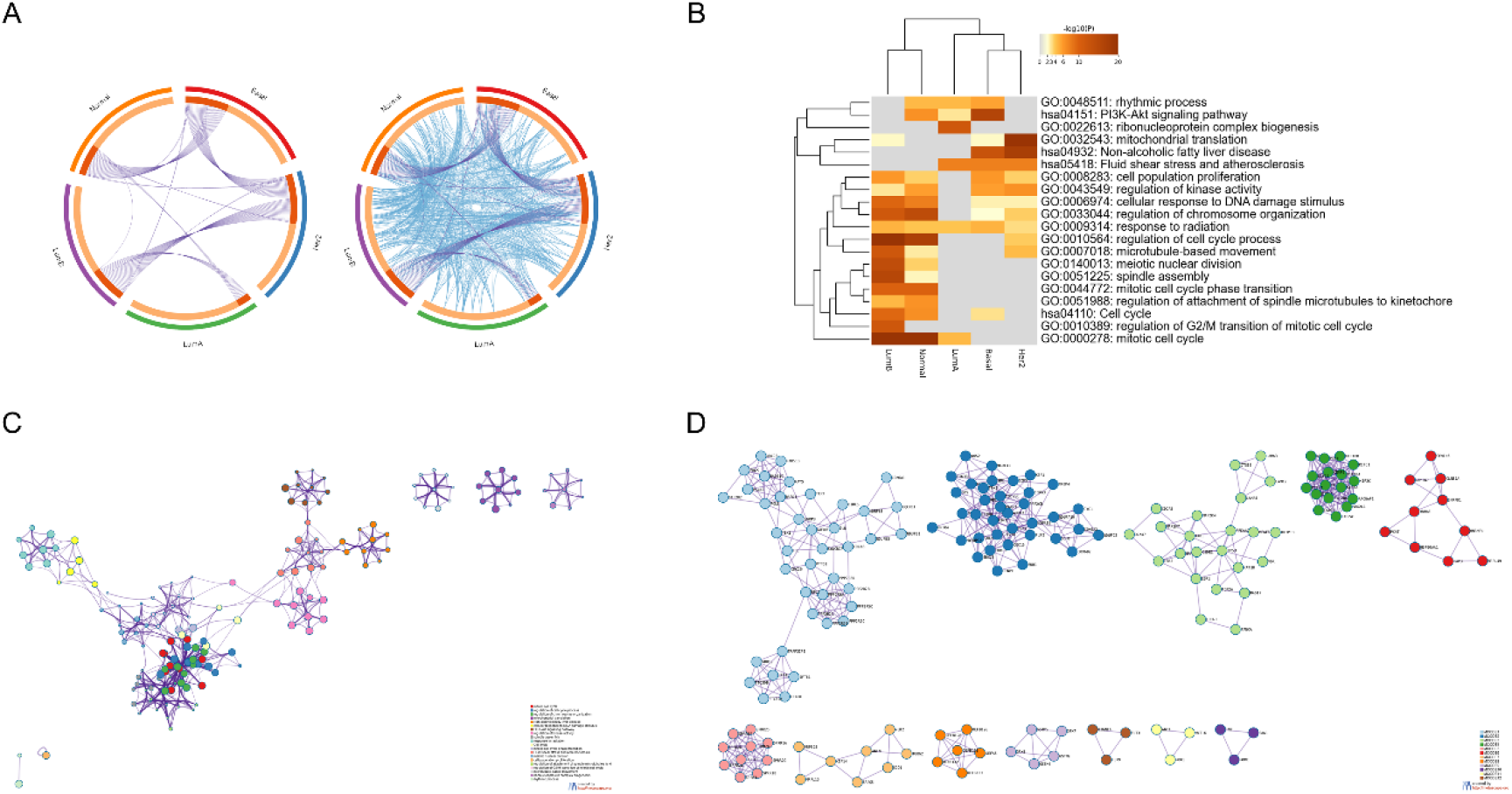
GO and KEGG enrichment analysis of top100 genes. **A**. Gene overlap analysis expanded via shared enriched ontologies. On the outside, each arc represents the identity of each gene list: LumA (green arc), LumB (purple arc), Normal (orange arc), Basal (red arc) and Her2 (blue arc). On the inside, each arc represents a gene list, where each gene member of that list is assigned a spot on the arc. Dark orange color represents the genes that are shared by multiple lists and light orange color represents genes that are unique to that gene list. Purple lines link the same gene that are shared by multiple gene lists (notice a gene that appears in two gene lists will be mapped once onto each gene list, therefore, the two positions are purple linked). Blue lines link the genes, although different, fall under the same ontology term (the term has to statistically significantly enriched and with size no larger than 100). **B**. For the four input gene lists, GO and KEGG enrichment terms were hierarchically clustered into a tree based on Kappa-statistical similarities among their gene memberships. The heatmap cells are colored by their P values, white cells indicate the lack of enrichment for that term in the corresponding gene list. **C**. Network layout of a subset of representative terms from the full cluster. Each term is represented by a circle node, where its size is proportional to the number of input genes fall under that term, and its color represent its cluster identity. Terms with a similarity score > 0.3 are linked by an edge (the thickness of the edge represents the similarity score). **D**. MCODE algorithm was then applied to this network to identify neighborhoods where proteins are densely connected. Network nodes are displayed as pies. Color code for pie sector represents a gene list and is consistent with the colors used in panel (A).

Since the model has the highest prediction precision for basal-like samples, we further explored the patient-specific feature pattern of 23 basal-like samples of the validation data set (Table S4). The basal-like group composes up to 15% of breast cancer and is characterized by triple-negative profile and is associated with an aggressive phenotype, high histological grade, poor clinical behavior, and high rates of recurrences and/or metastasis [44]. Among all frequently selected genes, CYP4X1, PPP2R5A, and SPRR2G are ranked in the top200 genes of all 23 basal-like samples. CYP4X1 encodes a member of the cytochrome P450 superfamily of enzymes. Cytochrome P450s are key oxidative enzymes that metabolize many carcinogens and anticancer drugs. A tissue microarray study found that multiple cytochrome P450s can be used as prognostic markers of breast cancer[45]. The product of PPP2R5A belongs to the phosphatase 2A(PP2A) regulatory subunit B family. PPP2R5A plays an important role in many cellular activities by regulating the cellular location, substrate specification, and protein phosphatase function of PP2A. PPP2R5A is reported to be related to many diseases including cancers. An elevated expression of cyclin G is observed in many cancers including breast cancer and cyclin G binds to PPP2R5A resulting in PP2A desphosphorylating MDM2, thus enhancing the degradation of P53[46]. SPRR2G (Small Proline-Rich Protein 2) is a member of SPRR family of cornified envelope precursor proteins. Besides, SPRR1A (22/23), SPRR2D (22/23), SPRR3 (22/23), and SPRR2A (22/23) were found in the top 200 genes of most samples. SPRR3 promotes breast cancer cell proliferation by enhancing p53 degradation via the AKT and MAPK pathways and its downregulation is associated with triple-negative breast cancer (TNBC), which is the main component of basal-like breast cancer[47]. SPRR family genes and FLNB are involved in the epidermal cell differentiation pathway, which is a specific enrichment pathway for basal-like samples. FLNB has the function of regulating EMT[48], and meanwhile, it’s reported that SPRR can play a role as a ligand of SH3 protein domain and is related to EMT in cholangiocarcinoma[49]. Basal-like samples are highly invasive and show EMT characteristics, but the association between FLNB and basal-like breast cancer needs to be further verified. In the top100 gene list (Table S4), tumor suppressor EP300 (22/23), estrogen receptor ESR2 (22/23), G protein-coupled estrogen receptor GPER1 (22/23), BACE1 (21/23), colony stimulating factor-1 receptor CSF1R (21/23) and formation of transcription factor complex AP-1 related FOS (21/23) are all reported to be linked with development and progression of breast cancer[50–53].

### Validation in unlabeled breast cancer samples

To explore the robustness and generalization performance of cAGCN, the remaining unlabeled 88 samples of TCGA BRCA were preprocessed for prediction. Most of them were predicted as luminal A (52/88, 59.10%) and luminal B (20/88, 22.73%), which was consistent with the distribution proportion of breast cancer subtypes. The predicted subgroups have a log-rank P value of 0.043 showing a significant survival difference in this small cohort (**Fig** S2).

## Discussion

Benefitting from the rapid advancement of omics technologies and existing large labeled biomedical data, more supervised methods are developed for patient diagnosis and biological knowledge discovery. High-dimensionality and small sample size, as well as the different distributions of omics data have posed great challenges to multi-omics integrative analysis. A well-designed architecture is needed which can jointly make accurate decisions and learn the interrelationships between various biological data types or entities. In this paper, we proposed a supervised deep learning architecture, AGCN, for breast cancer subtype identification. We utilized GCN for taking advantages of various data features and gene inherent structures and attention mechanism for flexibly learning the association patterns of different omics data types or biomolecules. Through ablation studies, both GCN and attention mechanism were proved to be essential to multi-omics integrative classification. The distribution of attention scores extracted from the two omics-specific attention mechanism suggests that gene expression plays a predominant role in the biomolecular subtype identification of breast cancer. The observation is further validated through performance comparison of different combinations of omics data types as lack of mRNA information significantly reduces the performance of cAGCN. Beyond this, exploiting complementary information of different molecular data can also improve the prediction of breast cancer molecular subtypes since cAGCN trained with three data types has achieved the best classification performance among all datasets.

Deep learning methods have been widely applied in many fields such as computer vision and natural language processing, however, deep models designed for biomedical data are often limited in practice for lack of interpretation. Appropriate interpretation methods can auxiliary in transforming black-box models into white-box models. Thus, LRP algorithm was applied to explain how each gene related to the model decision. Our findings demonstrated that LRP combined with a well-trained deep model could present meaningful patient-specific or subtype-specific factors. For basal-like breast cancer patients, biomarkers (e.g., CYP4X1, PPP2R5A, SPRR2G, ESR2, BACE1, EP300) were identified as the highly influential biomarkers and the various gene rankings across different patients also demonstrated the individual inherent heterogeneity.

Since we only focused on the highly variable features of different omics data types, such feature selection strategy may introduce bias and ignore those slightly variable but functionally indispensable biomarkers. Although manual addition of reference biomarkers can help alleviate the problem, it is not applicable to analysis of unknown diseases. Development of an end-to-end approach with no need of complicated preprocessing procedures can be one of the solutions.

## Materials and Methods

### TCGA BRCA dataset

We downloaded the TCGA BRCA benchmark dataset from[54], which included 759 samples with omics data of CNV at the genome level, DNA methylation, and mRNA expression at the transcriptome level. The real 671 patient labels came from the results of a previous TCGA cohort study[55] and the remaining 88 unknown samples were used for external validation. For mRNA expression and DNA methylation data, median absolute deviation (MAD) was adopted to select the top 2,000 most-variable features. For CNV data, genes in significantly gained or lost regions (q-value<0.05) reported in Broad Institute for breast cancer were selected. 3,167 high variable genes (hvGs) at different molecular levels were retained (**Fig 1A**) and the corresponding dataset was named as MDC. Besides, we selected genes from DisGenet[56] which were reported to be breast cancer related and filtered out genes with a GDA score < 0.2. Thus, a total of 540 genes were collected as reference gene sets and the corresponding dataset was named as MDC_addRef consisting of 3483 genes.

### Protein-protein interaction network

We downloaded human PPI from StringDB (v11.5) as the graph structure input, which consisted of 19,385 vertices and 11,938,498 edges. After removing low confidence edges (confidence score < 850) and ID mapping, the final reference PPI contained 12,397 vertices and 287,084 edges. Different gene sequencing platforms are always hampered by missing data due to discrepant gene annotations and only a minority of genes are disease-related. To reduce irrelevant features and further improve the quality of the input biological network, we mainly focused on 3,167 hvGs at different molecular levels. Considering that direct edge reduction would affect the relationships between the original gene pairs and result in the loss of association information, we further explored the performance of GCN with network input preserving neighborhoods of different order.

### Implement of GCN

Graph Convolutional Neural Networks define spectral convolution operations on graphs. Referring to the convolution theorem, the spectral space signal obtained by the Fourier transform is first linearly transformed and the inverse Fourier transform is then used to convert the spectral space signal to the original space to realize GCN. [57] simplify and normalize the convolution kernel of Chebyshev polynomial approximation to obtain a first-order GCN layer which is shown in Equation (1), where 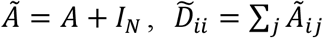, *W*^(*l*)^ is the trainable parameters of the *l* th layer, σ represents the activation function. Here, we used GCN to aggregate the information of gene neighboring nodes and then update the feature representation of genes.

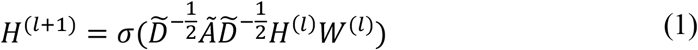

### Attention mechanism

Attention mechanism is a special module in machine learning model that can automatically learn and calculate the association between input data and output data. Here, we explored the influences of three attention mechanism modules on the model performance.

▪ Squeeze and Excitation

We referred to the proposed architecture which was used to explicitly model the interdependencies between channels of convolutional neural networks[58]. Squeeze and Excitation (SE) block here is in use for importance assessment of different omics-level features. Squeeze is the global information embedding module using global average pooling to get channel statistics. The purpose of Excitation is to capture the complex nonlinear interaction between channels, which is implemented through fully connected layers (FC) and gating mechanisms. Finally, residual connection is applied for output. The concrete structure is demonstrated in **Fig 5A** and the calculation steps are as follows:

**Fig 5.**
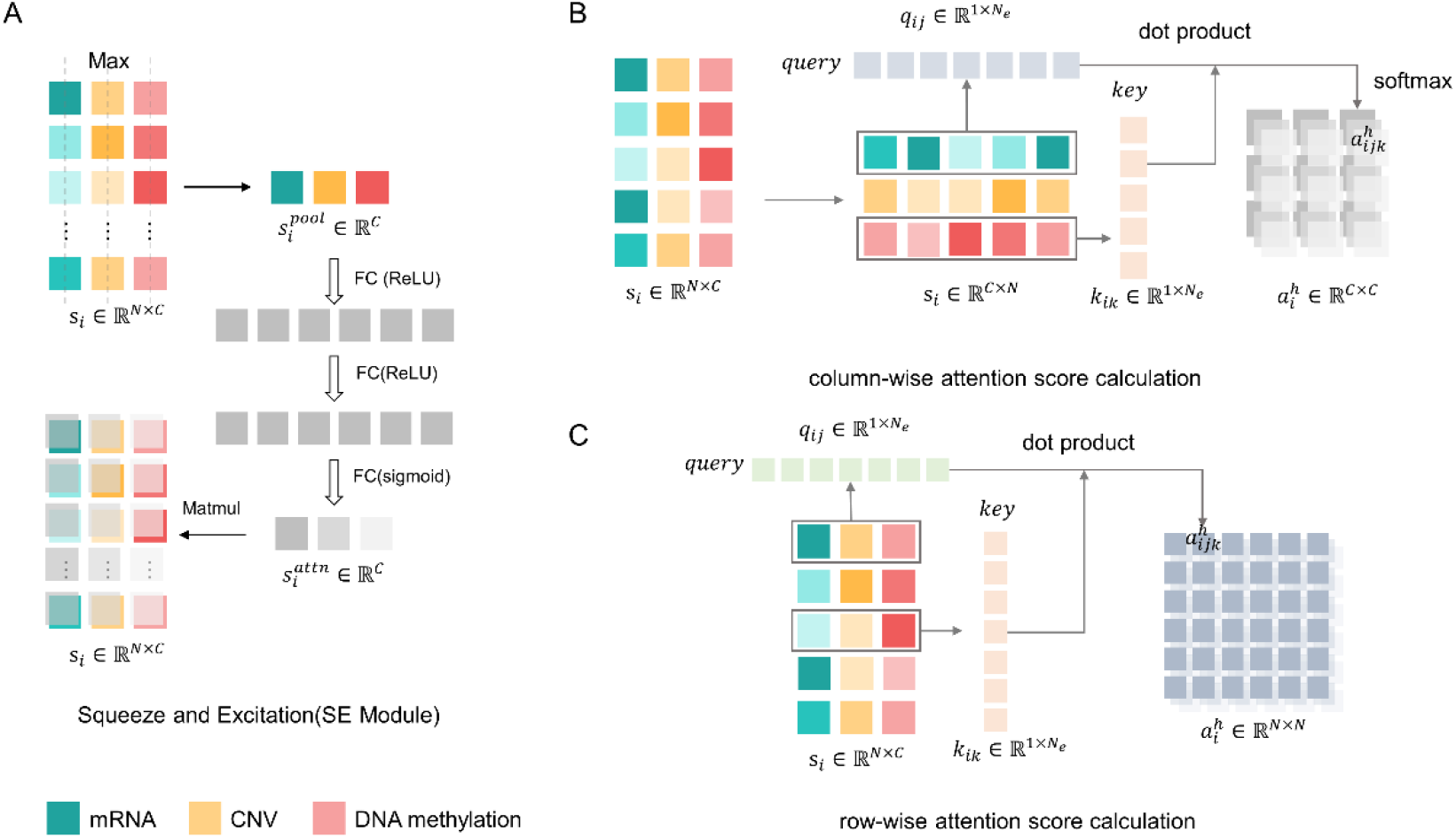
Implement of three attention mechanisms. **A**. Squeeze and Excitation (SE) module. **B**. Column-wise attention (cAttn). **C**. Row-wise attention (rAttn).

For sample *i, X*_*i*_ ∈ ℝ^*F*×*C*^,

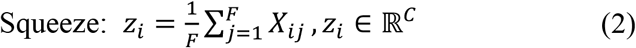

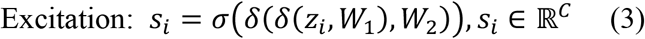

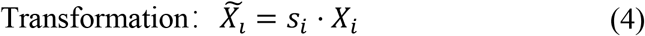

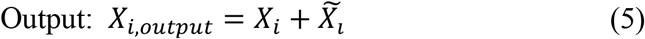

where δ represents the activation function ReLU and σ represents the activation function Sigmoid.

▪ column-wise self-attention and row-wise self-attention

The self-attention mechanism is a module that learns the internal associations of sequences. Referring to Scaled Dot-Product Attention[59], we first calculate the query (Q), key (K), and value (Q) vector corresponding to the input sequence, and calculate the dot product between the Q and K as the association evaluation between the two vectors and normalization is performed to obtain the final attention score (**Fig 5B, C**). Column-wise self-attention (cAttn) is used to capture the association between different omics and row-wise self-attention (rAttn) is used to capture association information between genes. The difference between the two is the implement of matrix transpose. GCN models with these three attention blocks were named as SEGCN (SE+GCN), cAGCN (cAttn+GCN), and rAGCN (rAttn + GCN) respectively.

### Layer-wise relevance propagation(LRP)[60]

The LRP algorithm utilizes the structure of the network by propagating the explanations from the output to the input layer based on the deep Taylor decomposition to explain the nonlinear decision. At some chosen root point 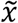, so that 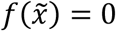, the first-order Taylor approximation of the function *f*(*x*) can be expressed as:

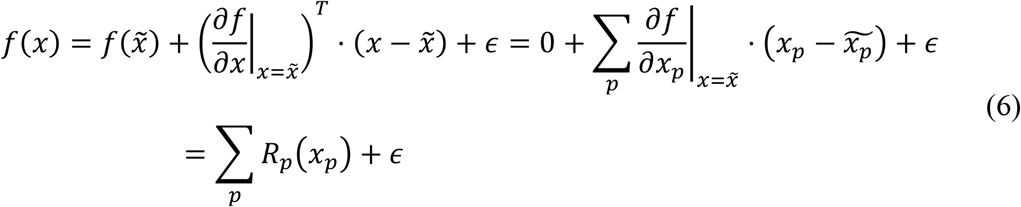

where *R*_*p*_ can represent the relevance between the input element *p* and the predicted output. For deep neural network layers with nonlinear activation function ReLU, of which the form is:

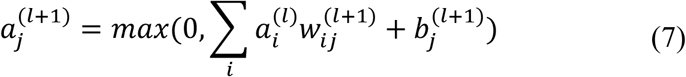

where 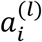 represents the output of neuron *i* of the *l* -th layer, 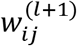 is the weight between the neuron *i* of the *l*-th layer and the neuron *j* of the (*l* + 1)-th layer and 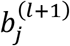 is bias. Since the layers of this type always have non-negative activations, the relevance propagation rule is made as following according to z^+^-rule:

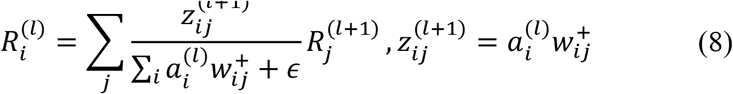

We supposed that the outputs of the global pooling layer are features at the gene level which have already integrated cross-omics relationships and gene structural information, named fusion feature. LRP was carried out on layers right after graph convolution layers to measure the contribution of genes with fusion feature to the model decision and LRP rule is not conducted on batch normalization layers.

Specially, we took the sample-average score as the relevance/contribution of a gene corresponding to subtype C:

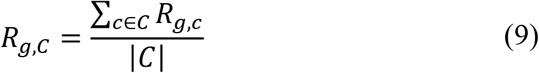

### Evaluation strategies and metrics

5-fold cross-validation was employed to evaluate the performance of all models on the TCGA BRCA datasets. The best model reported in cross-validation was chosen as the final model for the downstream analysis. In case of the multi-class prediction task, area under the curve (AUC), balanced accuracy (ACC), and Matthew correlation coefficient (MCC) were used to evaluate the overall performance of all models. The mean and standard deviation of the evaluation metrics across different models were reported.

### Model parameters

In ablation study, network hyperparameters were determined by cross-validation. Finally, we set the dimensions of feed forward network to *d*-512-256-5 for cAMLP, *d*-1024-512-5 for simple GCN, *d*-512-256-5 for SEGCN, *d*-512-256-5 for cAGCN and *d*-512-5 for rAGCN where *d* is the data-space dimension. The dimension of graph convolution layer was set as *d*-64 for simple GCN, *d*-32 for SEGCN, cAGCN and rAGCN. The dimension of attention embedded layer was set as *d*-32 for SE, *d*-256 for cAttn and *d* -64 for rAttn. Dropout was adopted to prevent network from overfitting and was set to 0.5 for simple GCN, SEGCN, cAGCN and rAGCN and 0.8 for cAMLP. In comparison performance of different datasets, network hyperparameters were fixed for all datasets. Specifically, we set *d*-512-256-5 for feed forward network, *d*-32 for graph convolution layer and *d*-256 for cAttn.

### Univariable Cox PH analysis and enrichment analysis

Univariable Cox PH regression analysis was performed using R package *Survival* to screen genes associated with breast cancer survival (log-rank P value < 0.05). Functional analysis and DO enrichment analysis of the screened genes were implemented in R package *ClusterProfile* [61] and ClueGO[62] plugin in Cytoscape was used to visualize KEGG network. GO and KEGG functional enrichment analysis for multiple gene lists was performed using MetaScape[63]. Minimal overlap of 3 genes, enrichment factor of 1.5, and p-value of 0.05 were used for filtering. The remaining significant terms were then hierarchically clustered into a tree based on Kappa-statistical similarities among their gene memberships. Then 0.3 kappa score was applied as the threshold to cast the tree into term clusters. MCODE algorithm was further applied to the merged network to identify neighborhoods where proteins are densely connected. Pathway and process enrichment analysis has been applied to each MCODE component independently, and the three best-scoring terms by P value have been retained as the functional description of the corresponding components.

## Data Availability

The TCGA BRCA multi-omics data used in this paper were downloaded from[54] and PPI data was downloaded from https://stringdb-static.org/.

## Code Availablility

Source code is also available at Github at https://github.com/cicGuoH/GCN_multiomics.

## Acknowledgements

This work was supported by the grants from the National Natural Science Foundation of China (no. 22173065).

## Authors’ contributions

LM and LY conceived the project. LY and GH designed the project. GH analyzed and interpreted the data and wrote the manuscript. LM, LY and LX revised and contributed to the final manuscript. All authors have read the manuscript and agreed to its publication.

## Reference

1. Perou CM, Sørlie T, Eisen MB, van de Rijn M, Jeffrey SS, Rees CA, et al. Molecular portraits of human breast tumours. Nature. 2000;406: 747–752. doi:10.1038/35021093

2. Hu Z, Fan C, Oh DS, Marron J, He X, Qaqish BF, et al. The molecular portraits of breast tumors are conserved across microarray platforms. BMC Genomics. 2006;7: 96. doi:10.1186/1471-2164-7-96

3. Sørlie T, Tibshirani R, Parker J, Hastie T, Marron JS, Nobel A, et al. Repeated observation of breast tumor subtypes in independent gene expression data sets. Proc Natl Acad Sci. 2003;100: 8418–8423. doi:10.1073/pnas.0932692100

4. The Cancer Genome Atlas Network. Comprehensive molecular portraits of human breast tumours. Nature. 2012;490: 61–70. doi:10.1038/nature11412

5. Parker JS, Mullins M, Cheang MCU, Leung S, Voduc D, Vickery T, et al. Supervised Risk Predictor of Breast Cancer Based on Intrinsic Subtypes. J Clin Oncol. 2009;27: 1160–1167. doi:10.1200/JCO.2008.18.1370

6. Cascianelli S, Molineris I, Isella C, Masseroli M, Medico E. Machine learning for RNA sequencing-based intrinsic subtyping of breast cancer. Sci Rep. 2020;10: 14071. doi:10.1038/s41598-020-70832-2

7. Pan X, Hu X, Zhang Y-H, Chen L, Zhu L, Wan S, et al. Identification of the copy number variant biomarkers for breast cancer subtypes. Mol Genet Genomics. 2019;294: 95–110. doi:10.1007/s00438-018-1488-4

8. Szyf M. DNA methylation signatures for breast cancer classification and prognosis. Genome Med. 2012;4: 26. doi:10.1186/gm325

9. Hasin Y, Seldin M, Lusis A. Multi-omics approaches to disease. Genome Biol. 2017;18: 83. doi:10.1186/s13059-017-1215-1

10. METABRIC Group, Curtis C, Shah SP, Chin S-F, Turashvili G, Rueda OM, et al. The genomic and transcriptomic architecture of 2,000 breast tumours reveals novel subgroups. Nature. 2012;486: 346–352. doi:10.1038/nature10983

11. Taskesen E, Babaei S, Reinders MM, de Ridder J. Integration of gene expression and DNA-methylation profiles improves molecular subtype classification in acute myeloid leukemia. BMC Bioinformatics. 2015;16: S5. doi:10.1186/1471-2105-16-S4-S5

12. Gao F, Wang W, Tan M, Zhu L, Zhang Y, Fessler E, et al. DeepCC: a novel deep learning-based framework for cancer molecular subtype classification. Oncogenesis. 2019;8: 44. doi:10.1038/s41389-019-0157-8

13. Zhang G, Peng Z, Yan C, Wang J, Luo J, Luo H. MultiGATAE: A Novel Cancer Subtype Identification Method Based on Multi-Omics and Attention Mechanism. Front Genet. 2022;13: 855629. doi:10.3389/fgene.2022.855629

14. Yang B, Zhang Y, Pang S, Shang X, Zhao X, Han M. Integrating Multi-Omic Data With Deep Subspace Fusion Clustering for Cancer Subtype Prediction. IEEE/ACM Trans Comput Biol Bioinform. 2021;18: 216–226. doi:10.1109/TCBB.2019.2951413

15. Althubaiti S, Kulmanov M, Liu Y, Gkoutos GV, Schofield P, Hoehndorf R. DeepMOCCA: A pan-cancer prognostic model identifies personalized prognostic markers through graph attention and multi-omics data integration. 2021 [cited 21 May 2022]. doi:10.1101/2021.03.02.433454

16. Schulte-Sasse R, Budach S, Hnisz D, Marsico A. Integration of multiomics data with graph convolutional networks to identify new cancer genes and their associated molecular mechanisms. Nat Mach Intell. 2021;3: 513–526. doi:10.1038/s42256-021-00325-y

17. Wang C, Zhao N, Sun K, Zhang Y. A Cancer Gene Module Mining Method Based on Bio-Network of Multi-Omics Gene Groups. Front Oncol. 2020;10: 1159. doi:10.3389/fonc.2020.01159

18. Napolitano F, Zhao Y, Moreira VM, Tagliaferri R, Kere J, D’Amato M, et al. Drug repositioning: a machine-learning approach through data integration. J Cheminformatics. 2013;5: 30. doi:10.1186/1758-2946-5-30

19. Subramanian I, Verma S, Kumar S, Jere A, Anamika K. Multi-omics Data Integration, Interpretation, and Its Application. Bioinforma Biol Insights. 2020;14: 1177932219899051. doi:10.1177/1177932219899051

20. Nicora G, Vitali F, Dagliati A, Geifman N, Bellazzi R. Integrated Multi-Omics Analyses in Oncology: A Review of Machine Learning Methods and Tools. Front Oncol. 2020;10: 1030. doi:10.3389/fonc.2020.01030

21. Lin Y, Zhang W, Cao H, Li G, Du W. Classifying Breast Cancer Subtypes Using Deep Neural Networks Based on Multi-Omics Data. Genes. 2020;11: 888. doi:10.3390/genes11080888

22. Chaudhary K, Poirion OB, Lu L, Garmire LX. Deep Learning–Based Multi-Omics Integration Robustly Predicts Survival in Liver Cancer. Clin Cancer Res. 2018;24: 1248–1259. doi:10.1158/1078-0432.CCR-17-0853

23. Yang H, Chen R, Li D, Wang Z. Subtype-GAN: a deep learning approach for integrative cancer subtyping of multi-omics data. Robinson P, editor. Bioinformatics. 2021;37: 2231–2237. doi:10.1093/bioinformatics/btab109

24. Ramazzotti D, Lal A, Wang B, Batzoglou S, Sidow A. Multi-omic tumor data reveal diversity of molecular mechanisms that correlate with survival. Nat Commun. 2018;9: 4453. doi:10.1038/s41467-018-06921-8

25. Wang T, Shao W, Huang Z, Tang H, Zhang J, Ding Z, et al. MOGONET integrates multi-omics data using graph convolutional networks allowing patient classification and biomarker identification. Nat Commun. 2021;12: 3445. doi:10.1038/s41467-021-23774-w

26. Li X, Ma J, Leng L, Han M, Li M, He F, et al. MoGCN: A Multi-Omics Integration Method Based on Graph Convolutional Network for Cancer Subtype Analysis. Front Genet. 2022;13: 806842. doi:10.3389/fgene.2022.806842

27. Peng W, Wang J, Cai J, Chen L, Li M, Wu F-X. Improving protein function prediction using domain and protein complexes in PPI networks. BMC Syst Biol. 2014;8: 35. doi:10.1186/1752-0509-8-35

28. Buphamalai P, Kokotovic T, Nagy V, Menche J. Network analysis reveals rare disease signatures across multiple levels of biological organization. Nat Commun. 2021;12: 6306. doi:10.1038/s41467-021-26674-1

29. Huang JK. Systematic Evaluation of Molecular Networks for Discovery of Disease Genes. Cell Syst. 2018;6: 484–495. doi:10.1016/j.cels.2018.03.001

30. Shi Q, Zhang C, Peng M, Yu X, Zeng T, Liu J, et al. Pattern fusion analysis by adaptive alignment of multiple heterogeneous omics data. Wren J, editor. Bioinformatics. 2017;33: 2706–2714. doi:10.1093/bioinformatics/btx176

31. Singh A, Shannon CP, Gautier B, Rohart F, Vacher M, Tebbutt SJ, et al. DIABLO: an integrative approach for identifying key molecular drivers from multi-omics assays. Bioinformatics. 2019;35: 3055–3062. doi:10.1093/bioinformatics/bty1054

32. Qiu C, Yu F, Su K, Zhao Q, Zhang L, Xu C, et al. Multi-omics Data Integration for Identifying Osteoporosis Biomarkers and Their Biological Interaction and Causal Mechanisms. iScience. 2020;23: 100847. doi:10.1016/j.isci.2020.100847

33. Niu Z, Zhong G, Yu H. A review on the attention mechanism of deep learning. Neurocomputing. 2021;452: 48–62. doi:10.1016/j.neucom.2021.03.091

34. Montavon G, Lapuschkin S, Binder A, Samek W, Müller K-R. Explaining nonlinear classification decisions with deep Taylor decomposition. Pattern Recognit. 2017;65: 211–222. doi:10.1016/j.patcog.2016.11.008

35. Samek W, Binder A, Montavon G, Lapuschkin S, Müller K-R. Evaluating the Visualization of What a Deep Neural Network Has Learned. IEEE Trans Neural Netw Learn Syst. 2017;28: 2660–2673. doi:10.1109/TNNLS.2016.2599820

36. Hanczar B, Zehraoui F, Issa T, Arles M. Biological interpretation of deep neural network for phenotype prediction based on gene expression. BMC Bioinformatics. 2020;21: 501. doi:10.1186/s12859-020-03836-4

37. Salim A, Sumitra S. Spectral Graph Convolutional Neural Networks in the Context of Regularization Theory. IEEE Trans Neural Netw Learn Syst. 2022; 1–12. doi:10.1109/TNNLS.2022.3177742

38. Park HW, Guan K-L. Regulation of the Hippo pathway and implications for anticancer drug development. Trends Pharmacol Sci. 2013;34: 581–589. doi:10.1016/j.tips.2013.08.006

39. Cruceriu D, Baldasici O, Balacescu O, Berindan-Neagoe I. The dual role of tumor necrosis factor-alpha (TNF-α) in breast cancer: molecular insights and therapeutic approaches. Cell Oncol. 2020;43: 1–18. doi:10.1007/s13402-019-00489-1

40. Polakis P. Wnt Signaling in Cancer. Cold Spring Harb Perspect Biol. 2012;4: a008052–a008052. doi:10.1101/cshperspect.a008052

41. Kaminska B, Wesolowska A, Danilkiewicz M. TGF beta signalling and its role in tumour pathogenesis. Acta Biochim Pol. 2005;52: 329–337. doi:10.18388/abp.2005_3446

42. Buck MB, Knabbe C. TGF-Beta Signaling in Breast Cancer. Ann N Y Acad Sci. 2006;1089: 119–126. doi:10.1196/annals.1386.024

43. de Kruijf EM, Dekker TJA, Hawinkels LJAC, Putter H, Smit VTHBM, Kroep JR, et al. The prognostic role of TGF-β signaling pathway in breast cancer patients. Ann Oncol. 2013;24: 384–390. doi:10.1093/annonc/mds333

44. Valentin MD, da Silva SD, Privat M, Alaoui-Jamali M, Bignon Y-J. Molecular insights on basal-like breast cancer. Breast Cancer Res Treat. 2012;134: 21–30. doi:10.1007/s10549-011-1934-z

45. Murray GI, Patimalla S, Stewart KN, Miller ID, Heys SD. Profiling the expression of cytochrome P450 in breast cancer: Cytochrome P450 and breast cancer. Histopathology. 2010;57: 202–211. doi:10.1111/j.1365-2559.2010.03606.x

46. Mao Z, Liu C, Lin X, Sun B, Su C. PPP2R5A: A multirole protein phosphatase subunit in regulating cancer development. Cancer Lett. 2018;414: 222–229. doi:10.1016/j.canlet.2017.11.024

47. Kim JC, Yu JH, Cho YK, Jung CS, Ahn SH, Gong G, et al. Expression of SPRR3 is associated with tumor cell proliferation in less advanced stages of breast cancer. Breast Cancer Res Treat. 2012;133: 909–916. doi:10.1007/s10549-011-1868-5

48. Li J, Choi PS, Chaffer CL, Labella K, Hwang JH, Giacomelli AO, et al. An alternative splicing switch in FLNB promotes the mesenchymal cell state in human breast cancer. eLife. 2018;7: e37184. doi:10.7554/eLife.37184

49. Demetris AJ, Specht S, Nozaki I, Lunz JG, Stolz DB, Murase N, et al. Small proline-rich proteins (SPRR) function as SH3 domain ligands, increase resistance to injury and are associated with epithelial–mesenchymal transition (EMT) in cholangiocytes. J Hepatol. 2008;48: 276–288. doi:10.1016/j.jhep.2007.09.019

50. Asaduzzaman M, Constantinou S, Min H, Gallon J, Lin M-L, Singh P, et al. Tumour suppressor EP300, a modulator of paclitaxel resistance and stemness, is downregulated in metaplastic breast cancer. Breast Cancer Res Treat. 2017;163: 461–474. doi:10.1007/s10549-017-4202-z

51. Molina L, Bustamante FA, Bhoola KD, Figueroa CD, Ehrenfeld P. Possible role of phytoestrogens in breast cancer via GPER-1/GPR30 signaling. Clin Sci. 2018;132: 2583–2598. doi:10.1042/CS20180885

52. Patsialou A, Wang Y, Pignatelli J, Chen X, Entenberg D, Oktay M, et al. Autocrine CSF1R signaling mediates switching between invasion and proliferation downstream of TGFβ in claudin-low breast tumor cells. Oncogene. 2015;34: 2721–2731. doi:10.1038/onc.2014.226

53. Langer S, Singer CF, Hudelist G, Dampier B, Kaserer K, Vinatzer U, et al. Jun and Fos family protein expression in human breast cancer: correlation of protein expression and clinicopathological parameters. Eur J Gynaecol Oncol. 2006;27: 345–352.

54. Duan R, Gao L, Gao Y, Hu Y, Xu H, Huang M, et al. Evaluation and comparison of multi-omics data integration methods for cancer subtyping. Wang E, editor. PLOS Comput Biol. 2021;17: e1009224. doi:10.1371/journal.pcbi.1009224

55. Sanchez-Vega F, Mina M, Armenia J, Chatila WK, Luna A, La KC, et al. Oncogenic Signaling Pathways in The Cancer Genome Atlas. Cell. 2018;173: 321-337.e10. doi:10.1016/j.cell.2018.03.035

56. Piñero J, Bravo À, Queralt-Rosinach N, Gutiérrez-Sacristán A, Deu-Pons J, Centeno E, et al. DisGeNET: a comprehensive platform integrating information on human disease-associated genes and variants. Nucleic Acids Res. 2017;45: D833–D839. doi:10.1093/nar/gkw943

57. Kipf TN, Welling M. Semi-Supervised Classification with Graph Convolutional Networks. 2017. doi:10.48550/arXiv.1609.02907

58. Jie H, Shen L, Sun G. Squeeze-and-Excitation Networks. 2018. pp. 7132–7141. Available: https://openaccess.thecvf.com/content_cvpr_2018/papers/Hu_Squeeze-and-Excitation_Networks_CVPR_2018_paper.pdf

59. Vaswani A, Shazeer N, Parmar N, Uszkoreit J, Jones L, Gomez AN, et al. Attention is All you Need. Advances in Neural Information Processing Systems. Curran Associates, Inc.; 2017. Available: https://proceedings.neurips.cc/paper/2017/hash/3f5ee243547dee91fbd053c1c4a845aa-Abstract.html

60. Bach S, Binder A, Montavon G, Klauschen F, Müller K-R, Samek W. On Pixel-Wise Explanations for Non-Linear Classifier Decisions by Layer-Wise Relevance Propagation. Suarez OD, editor. PLOS ONE. 2015;10: e0130140. doi:10.1371/journal.pone.0130140

61. Yu G, Wang L-G, Han Y, He Q-Y. clusterProfiler: an R Package for Comparing Biological Themes Among Gene Clusters. OMICS J Integr Biol. 2012;16: 284–287. doi:10.1089/omi.2011.0118

62. Bindea G, Mlecnik B, Hackl H, Charoentong P, Tosolini M, Kirilovsky A, et al. ClueGO: a Cytoscape plug-in to decipher functionally grouped gene ontology and pathway annotation networks. Bioinformatics. 2009;25: 1091–1093. doi:10.1093/bioinformatics/btp101

63. Zhou Y, Zhou B, Pache L, Chang M, Khodabakhshi AH, Tanaseichuk O, et al. Metascape provides a biologist-oriented resource for the analysis of systems-level datasets. Nat Commun. 2019;10: 1523. doi:10.1038/s41467-019-09234-6

